# *Cntnap2* deletion in mice causes developmentally transient alterations in auditory processing and persistent startle hyperreactivity

**DOI:** 10.64898/2026.09.17.752490

**Authors:** N. Chan, S. D. I. Seheult, A. Warr, P. A. Faure, S. Schmid, K.Y. Choe

**Affiliations:** Department of Psychology, Neuroscience & Behaviour, McMaster University, Hamilton, Ontario, L8S 4K1, Canada; Department of Anatomy and Cell Biology, University of Western Ontario, London, Ontario, N6A 3K7, Canada

**Author notes:** Corresponding Author: Katrina Choe, Department of Psychology, Neuroscience & Behaviour, McMaster University, Hamilton, Ontario, Canada, L8S 4K1.

## Abstract

Autism spectrum disorder (ASD) is a neurodevelopmental condition often associated with auditory symptoms, including impairments in sound perception and hypersensitivity. These symptoms are thought to be partly due to the delayed development of the auditory brainstem, as evidenced through aberrant auditory brainstem responses (ABRs) and exaggerated startle responses in infants diagnosed with ASD. Loss-of-function mutations in *CNTNAP2*, a gene implicated in auditory processing and language development, are causally associated with ASD. Accordingly, knocking out *Cntnap2* causes auditory brainstem impairments in rats; however, it is unknown if similar disruptions are observed in mice due to a possible conserved role of *Cntnap2* in the auditory brainstem. To test this, we characterized the development of auditory processing, reactivity, and sensory filtering in juvenile and adult *Cntnap2* knockout (KO) mice. We found that juvenile KO mice exhibited larger ABR wave III amplitudes and smaller wave IV widths, which normalized in adult mice, suggesting the possibility of transient hyperexcitability in the developing auditory brainstem. Additionally, our acoustic startle measurements revealed sex-specific phenotypes in which male KO mice exhibit an exaggerated startle response that persists into adulthood, while prepulse inhibition (PPI) remained generally intact in KO mice at both ages. These results demonstrate that loss of *Cntnap2* in mice produces auditory brainstem abnormalities similar to rats, but causes species- and sex-specific effects on auditory reactivity, establishing the mouse model as a complementary animal model for investigating neural mechanisms underlying auditory dysfunction in ASD.

**Significance Statement:** Impairments in auditory processing are commonly reported amongst individuals diagnosed with autism spectrum disorder, but it is unclear how genetic risk factors can alter the brain circuits that underlie these auditory symptoms. Here, we used a mouse model with a high-confidence risk gene (*CNTNAP2*) to investigate how early auditory processing is impacted throughout development through both physiological and behavioural measures. We show that early auditory dysfunction is a conserved consequence of *Cntnap2* loss early in development but normalized in adulthood, similar to the *Cntnap2* knockout rat model. In contrast to the transient auditory brainstem response phenotype, we observed a sex-dependent alteration in auditory-evoked startle behaviour that persisted into adulthood.

This work highlights the importance of investigating how altered sensory processing may contribute to the development of ASD-related behaviours and reinforces the need to consider sex as a biological variable when investigating neural mechanisms underlying neurodevelopmental disorders.

## Introduction

The neural circuits that underlie sensory processing undergo extensive refinement during early development, and such changes are shaped by both genetics and environmental factors. This developmental process is thought to be disrupted in autism spectrum disorder (ASD), as many individuals diagnosed with ASD experience atypical sensory processing^1–4^. Among the most commonly reported sensory disruptions is atypical auditory processing, such as hypo- or hypersensitivity to sound, that often appear early in life^5,6^. Considering that auditory information is essential for many facets of daily life, including language acquisition and social communication, disruptions in early auditory processing can contribute to multiple behavioural features associated with ASD.

To investigate the neural basis behind these auditory symptoms, clinical studies have utilized electrophysiological measures to assess auditory processing in individuals with ASD. One of the most used techniques are auditory brainstem response (ABR) recordings, a non-invasive electrophysiological technique that measures the synchronized neural activity along the ascending auditory pathway^7^. Using this technique, studies have discovered that the initial encoding of auditory information within the auditory brainstem is disrupted in ASD - infants that are later diagnosed with ASD frequently display prolonged ABR wave latencies^8–11^ and increased wave amplitudes^12^, suggestive of disruptions of the neural responsivity along the ascending auditory pathway. Because the auditory brainstem projects to diverse brain regions, disruptions early in development may have lasting impacts on other related sensory and behavioural functions^13–16^. For instance, individuals with ASD commonly exhibit an exaggerated acoustic startle response^17,18^ and reduced prepulse inhibition (PPI) of acoustic startle, indicating diminished sensory filtering and sensorimotor gating of auditory inputs into the brainstem^19,20^. Beyond sensory-related symptoms, auditory processing abnormalities have also been associated with broader social and behavioural impairments in individuals with ASD^21–23^. For instance, auditory processing differences in infants was predictive of difficulties with adaptive behaviours^23^ and lower verbal intelligence quotient in school-aged children^21^. Together, these observations highlight the potential developmental significance of early sensory dysfunction and raise the possibility that it may contribute to the emergence of later behavioural phenotypes in ASD.

The genetic contribution to the aetiology of ASD provides an important framework for investigating the neural mechanisms that contribute to the auditory phenotypes^24^. Among the identified risk genes, both rare, high-impact loss-of function variants and more common genetic variants of the contactin-associated protein-like 2 (*CNTNAP2*) gene have been well-linked with ASD^25,26^. The *CNTNAP2* gene encodes for a cell adhesion protein known as CASPR2, which plays important roles in axonal organization and synapse development^27–29^. *CNTNAP2*/CASPR2 is highly expressed in auditory pathways, including early in development^27,30,31^. *CNTNAP2* mutations are associated with auditory dysfunction in humans^32,33^.

Similarly, *Cntnap2* knockout (KO) rats, which feature a complete loss of function mutation, exhibit several phenotypes consistent with a presence of auditory dysfunction; we have previously demonstrated that juvenile *Cntnap2* KO rats have lower amplitudes and longer ABR waveform latencies, indicative of the delayed development in the auditory brainstem^34–37^. Additionally, we have also shown that *Cntnap2* KO rats exhibit heightened acoustic startle and reduced PPI^34,35,38,39,37,40^, features commonly reported in individuals with ASD^17–20^. While these demonstrations made in *Cntnap2* KO rats are supportive of the potentially conserved role of *CNTNAP2* in auditory processing, continued mechanistic work could benefit from incorporating the *Cntnap2* KO mouse model, which would enable the use of a wide array of available mouse genetic tools^41,42^. Furthermore, with the exception of auditory phenotypes, the *Cntnap2* KO mouse model has been comprehensively characterized across cellular, circuit, and behavioural domains^43–52^, providing a well-established framework for determining how auditory dysfunction relates to broader neurobiological and behavioural alterations caused by the *Cntnap2* deletion. Notably, the first and only direct demonstration of auditory processing alterations in *Cntnap2* KO mice identified impairments in temporal sound processing associated with morphological differences within the medial geniculate nucleus in auditory thalamus^53^. However, a comprehensive characterization of auditory processing phenotypes across development remains lacking in *Cntnap2* KO mice.

Here, we characterized the developmental trajectory of auditory brainstem processing in *Cntnap2* KO mice. We recorded ABRs at two developmental stages - juveniles and adults - to assess hearing sensitivity, neural responsivity, and speed of neurotransmission in *Cntnap2* KO mice and wildtype (WT) littermates. Furthermore, we examined behavioural measures of auditory brainstem function by measuring the acoustic startle and PPI thresholds to determine if reactivity and sensorimotor gating (PPI) were affected in the *Cntnap2* KO mouse model. We found that juvenile *Cntnap2* KO mice exhibit altered neural responsivity in selective regions along the ascending auditory pathway that is normalized in adulthood. Additionally, juvenile *Cntnap2* KO mice displayed exaggerated acoustic startle responses that persisted in a sex-dependent manner in male KO animals, while the PPI response appeared to remain mostly intact in KO mice throughout development.

## Methods

### Animals

All experimental procedures were approved by the Animal Research Ethics Board of McMaster University and conformed to the *Guide to the Care and Use of Experimental Animals* published by the Canadian Council of Animal Care. The *Cntnap2* KO mice were obtained from the Jackson laboratories (stock #017482, C57BL/6J background strain) and bred in-house for 10+ generations. Heterozygous *Cntnap2* KO males and females were crossed to produce homozygous *Cntnap2* KO mice along with wildtype (WT) littermates. Both males and females were used in this study. Mice were housed in groups of 2 to 4 sex-matched littermates in polycarbonate cages (28 cm × 17.5 cm × 12 cm) with bedding, nesting material, and an opaque plastic tube for hiding and enrichment. All experimental mice were kept in a holding room with a 12 h light:12 h dark cycle (lights on at 08:00, lights off at 20:00) and had ad libitum access to food and water.

### Auditory Brainstem Recordings (ABRs)

ABRs were collected from age-separate cohorts between postnatal day (PND) 30-40 for juveniles and PND 56-70 in adults with the following sample sizes: WT juvenile (n = 13; 7 males/6 females), KO juvenile (n = 14; 5 males/9 females), WT adult (n = 12; 9 males/3 females), and KO adult (n = 8; 4 males/4 females). Mice were anesthetized with an intraperitoneal injection of ketamine/xylazine cocktail (0.1 to 0.25 ml of a 3:1 v/v mixture of 100 mg/ml Ketamine + 20 mg/ml Xylazine; final animal dose 3.2 to 8.0 mg/kg). Once sedated, they were placed on a foam pad inside an angle-iron Faraday cage (50 × 40 × 60 cm), fitted with sound-attenuating foam (Sonex® Classic; Pinta Acoustic, U.S.A.) backed by grounded copper meshing. Subdermal needle electrodes (3 lead disposable, 27-gauge; S83018–19 Rochester ElectroMedical, Lutz, Florida, USA) were inserted at three different locations: (1) the reference electrode at the nape of the neck, (2) the recording electrode placed at the midline axis between the two ears, and (3) the ground electrode placed at the hip of the mouse (**Fig. 1A**). Recorded ABR waveforms were amplified 20× by a low impedance head stage (TDT RA4LI) whose output was further amplified 250× and digitized by a preamplifier (TDT RA4PA Medusa) before passing to a multifunction processor (TDT RX6) via a fiber optic cable. Both the head stage and preamplifier were battery operated and located inside the Faraday cage.

**Figure 1.**
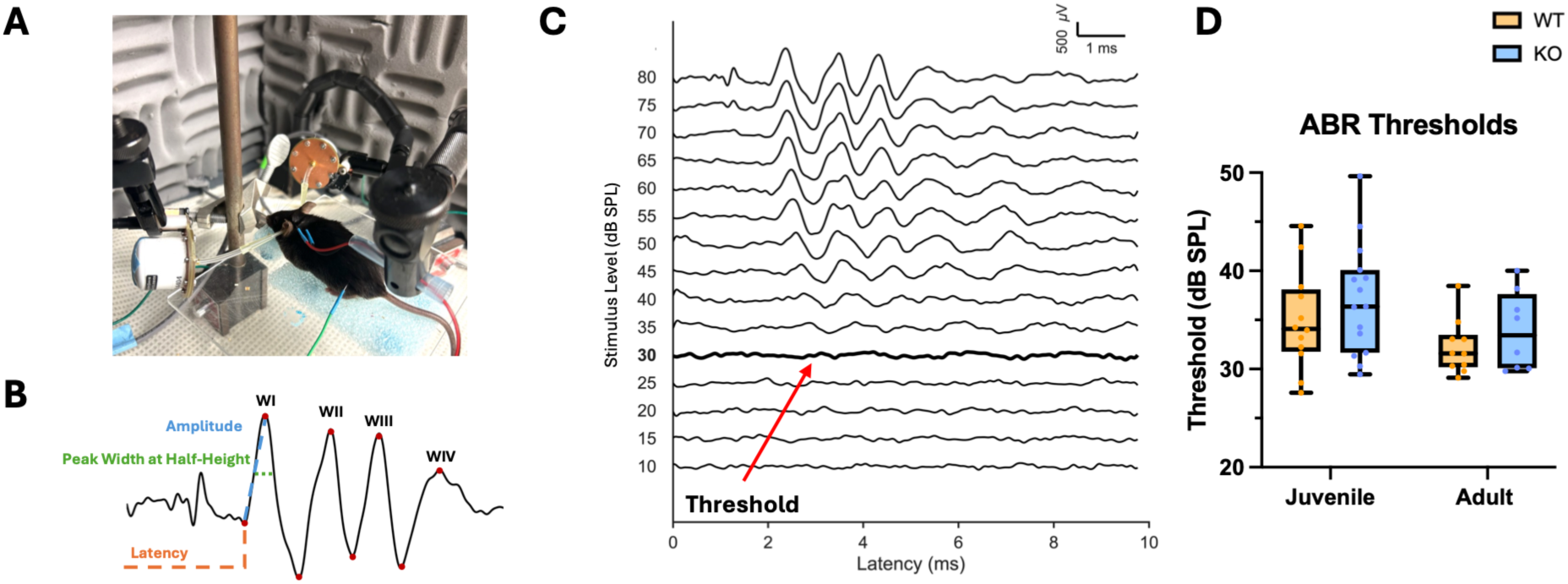
*Cntnap2* KO mice have normal auditory sensitivity with a typical maturation pattern. **(A)** Auditory brainstem response (ABR) experimental setup with an anesthetized adult mouse. Three subdermal leads are placed and speakers are positioned in the ear canal. **(B)** Example of an ABR trace from a WT mouse in response to 80 dB sound pressure level (SPL) click stimulus, reflecting the synchronized neural activity in the early ascending auditory pathway. Dotted lines show how peak-to- peak amplitude (blue), peak latency (orange), and peak width at half height (green) were determined. **(C)** Example of a stacked ABR waveform plot used to visually determine auditory thresholds (lowest intensity at which waves are still present), indicated by the red arrow and the thicker waveform. **(D)** ABR thresholds of *Cntnap2* KO mice and WT littermates at juvenile and adult stages. *Cntnap2* KO mice displayed a maturation pattern comparable to WT. *p < 0.05.

**Figure 2.**
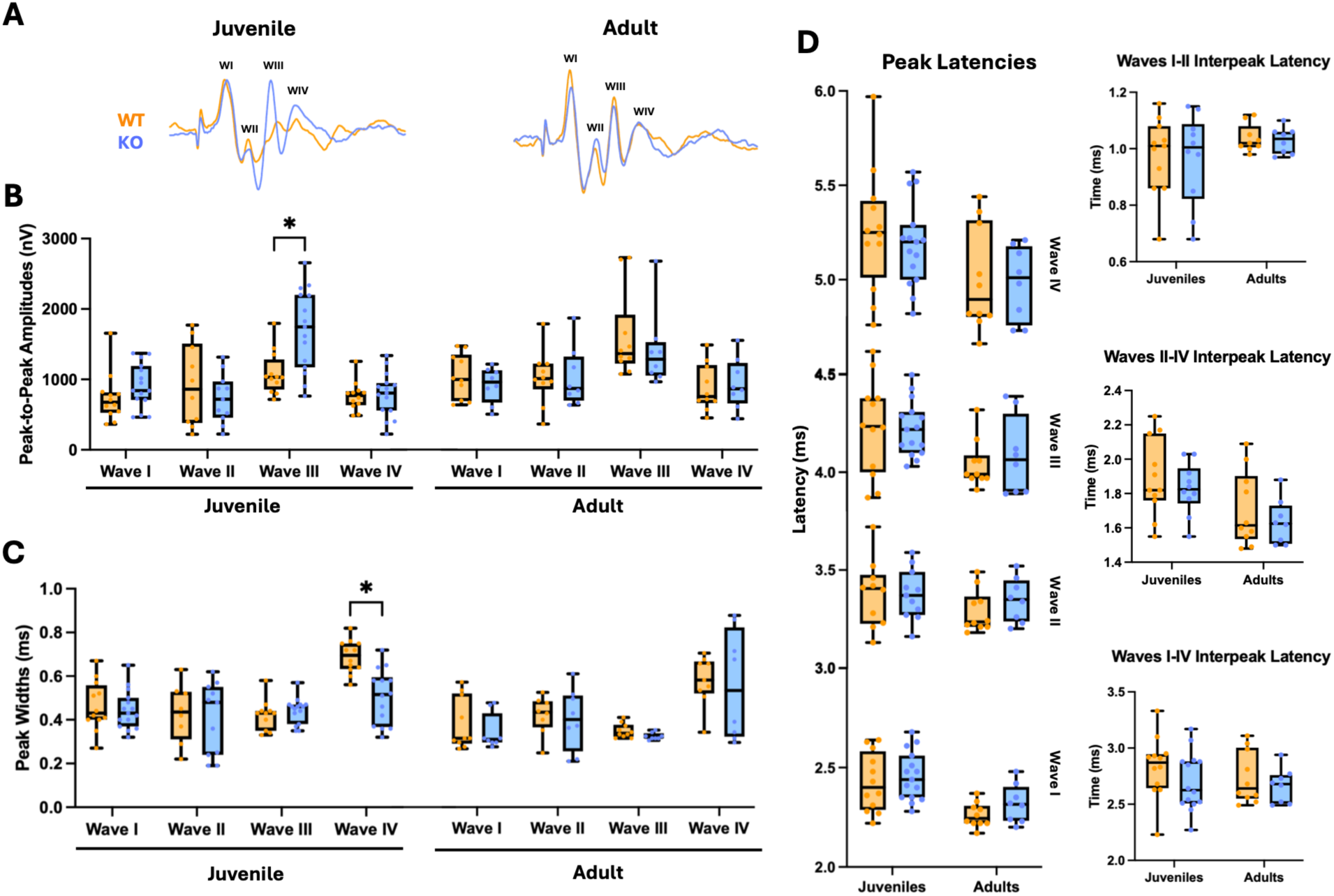
Juvenile *Cntnap2* KO mice show altered ABR waveforms that normalize in adults. **(A)** *Top*, Raw averaged ABR waveforms recorded from juvenile and adult WT (yellow) and *Cntnap2* KO (blue) mice to an 80 dB SPL click stimulus. ABR peak amplitudes shown as box and whisker plots displaying summary data for **B)** Peak-to-peak (P2P) amplitudes **C)** Peak widths at half height, and **D)** Peak latencies. \**p* < 0.05, \*\**p* < 0.01

The mice were exposed to repeated presentations of broadband acoustic clicks (duration= 1.0 ms, n = 512 repeats at 21 Hz) at 80 decibels sound pressure level (dB SPL re: 20 µPa) down to 10 dB SPL in 5 dB SPL steps. ABR waveforms were collected (recording window = 10 ms), averaged, then bandpass filtered (high-pass cut-off frequency = 300 Hz, low-pass cut-off frequency = 3 kHz, notch frequency = 60 Hz) before exporting for visualization and analysis. Upon completion of a recording, the electrodes were removed, cleaned with 95% ethanol, and the mouse was placed on a heating pad until it recovered. Mice were returned to their home cage once they showed full signs of wakeful behaviour.

To determine auditory thresholds (i.e. the lowest dB SPL still evokes a clear ABR response), ABR waveforms from individual mice were arranged by descending dB SPL and inspected visually by two blinded observers (**Fig. 1C**). The visually-inspected thresholds were then correlated with a MATLAB algorithm^54,55^ that applies a normalized cross-covariation analyses to output thresholds for varying criterion levels to standardize threshold estimates. Using this algorithm, we determined the auditory threshold by setting an optimal criterion value that correlated best to the visually determined thresholds which allowed for the standardization of threshold estimates across individual stacked plot displays.

ABR peak amplitudes have been used to measure neural responsivity across the auditory brainstem^56^. Each wave within the resultant ABR waveform have been shown to represent activity in specific structures along the ascending auditory pathway: auditory nerve (wave I), cochlear nucleus (CN; wave II), superior olivary complex (SOC; wave III), and inferior colliculus (IC; wave IV)^57^ (**Fig. 1B**). The peak-to-peak (P2P) amplitudes and latencies of peaks I-IV were determined using a bandpass filtered (cutoff frequencies 100 Hz and 1500 Hz) ABR waveform generated at 80 dB SPL for each animal.

Positive waveform peaks were identified using MATLAB’s *findpeaks* function. For each detected peak, the local minima (trough) preceding and following the peak was identified to calculate P2P amplitude (from trough to peak) and peak width at half height for manually labelled waves I-IV based on latency characteristics (**Fig. 1B**).

### Acoustic Startle Response and Prepulse Inhibition (PPI)

A separate cohort of mice underwent acoustic startle and PPI behavioural tests, assessed at both juvenile and adult timepoints (WT n = 21; 11 males/10 females, KO n = 19; 9 males/10 females). Mice were placed in an open, perforated, non-restricted plexiglass tube on a weight-transducing platform inside San Diego Instruments startle box system (SR-Lab System #2325-0400). Prior to testing, mice were handled for 5 minutes for three consecutive days by the experimenter. Additionally, on testing days, mice acclimated in the startle tube for 10 minutes with a background sound of 60 dB SPL white noise prior to testing conditions.

Mice underwent 3 consecutive days of testing. On the first day, a startle reactivity input/output (I/O) function was determined by presenting 11 different startle stimuli every 20 seconds presented in a pseudorandomized fashion (between 65-115 dB SPL white noise; 5 dB SPL increments, duration = 20 ms) on top of 60 dB white background noise. Each intensity level was tested 10 times, making a total of 110 trials, and the magnitude of the startle response was averaged at each intensity to determine the mean response across trials.

Startle magnitudes were defined as the maximum amplitude of the response waveform (**Fig. 3A**). The startle reactivity across the range of startle intensities was assessed by fitting each animal’s response to a sigmoidal regression curve in GraphPad Prism 11.0.0 (Nonlinear regression; Method: Sigmoidal, 4PL, X is concentration; Method: Least squares regression; Initial values: choose automatically; Confidence: Unstable parameters and ambiguous fits as Neither option; Diagnostics: default values including Adjusted R Squared, RMSE, and tests of normality, see ^58^):

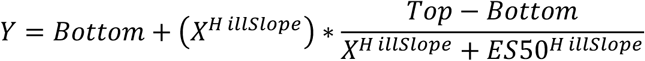

**Figure 3.**
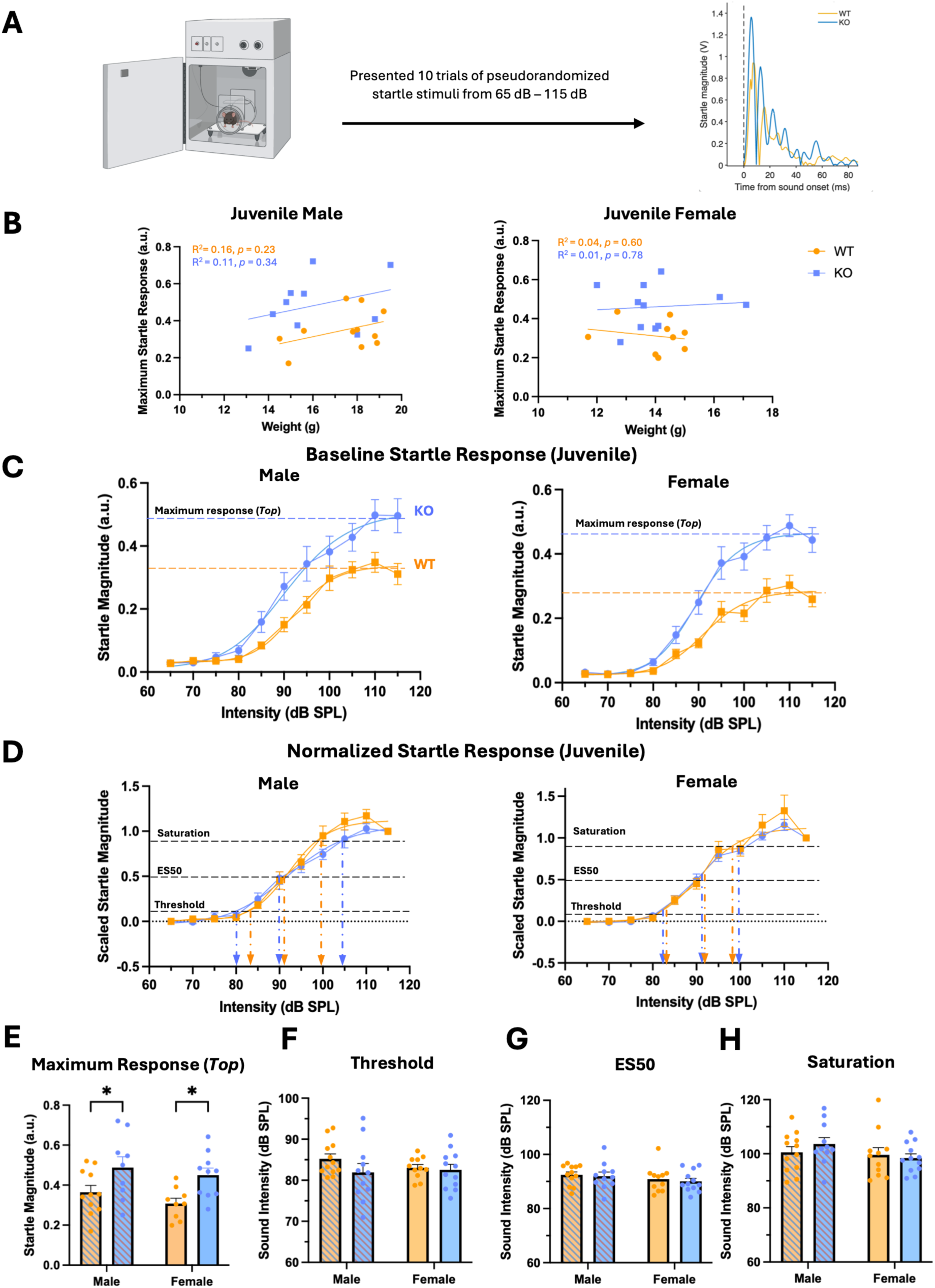
Male and Female juvenile *Cntnap2* KO mice show an exaggerated startle response. (A) Experimental timeline of the startle protocol and example trace from one juvenile WT and KO. **(B)** Correlation between weight and maximum startle response for each sex. Shown are I/O functions that represent the mean and standard error of mean (SEM) for each intensity with fitted sigmoidal curve for **(C)** Juvenile baseline startle response curves separated by sex**. (D)** Scaled startle response curve for each sex. Bar graphs show the mean, individual values, and error bars representing SEM for **(E)** Maximum startle response (*Top*), **(F)** Startle thresholds, **(G)** ES50, and **(H)** saturation points separated by sex. \**p* < 0.05

where *Y* represents the startle magnitude, *Bottom* is the minimum response on Y axis, *Top* is the maximum response magnitude, *X* is the intensity (dB SPL) needed to elicit the Y response, *ES50* is the intensity required to maintain half of the maximum response, and *Hillslope* is the slope of the sigmoidal curve. From this equation, parameters of interest were calculated and compared to assess differences in the baseline startle response, including the maximum startle response (Top), startle threshold (10% of maximum threshold response), ES50, and saturation point (90% of maximum startle response). In this sigmoidal regression analysis, GraphPad Prism provided the standard error of regression using Sy.x, which serves as an estimate of the goodness-of-fit for models involving two or more parameters.

Next, the response curve was scaled between 0 (response at the lowest intensity; 65 dB SPL) and 1 (response at the highest intensity; 115 dB SPL) to evaluate sound scaling. For each intensity level (X), the following equation was used to obtain scaled values:

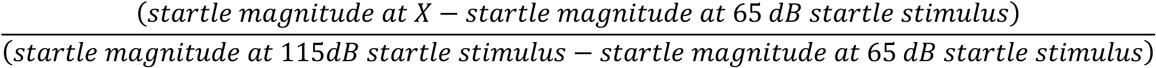

The scaled values were fit to a sigmoidal regression similar to above, with constrains for *Bottom* and *Top* set to 0 and 1 respectively (**Fig. 3D,H**; see also ^39^). ES50 was provided by the sigmoid regression, whereas the threshold and saturation point were calculated in MATLAB R2022a to solve for X:

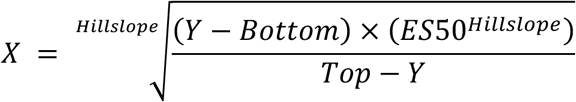

On the second and third day, we used a PPI paradigm to measure the decrease in startle amplitude of each mouse when a prepulse preceded a startle stimulus^34^. Mice were acclimated to the startle box for 10 minutes, then presented with a stimulus sequence as follows: 10 startle stimuli at 110 dB SPL, then 40 trials of randomized PPI conditions (75 or 85 dB SPL prepulse with an interstimulus interval, ISI, of either 30 or 100 ms). Following the PPI trials, another 10 trials of the startle stimulus alone were presented. All 20 startle alone trials were used to measure the baseline startle magnitude.

For PPI, the percent PPI was calculated with the following formula:

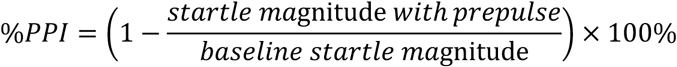

### Statistical Analysis

Statistical analyses were performed in GraphPad Prism 11.0.0 and RStudio Version 2026.01.0+392. The ABR data were presented as box and whisker plots, while startle thresholds were presented as bar graphs displaying the group mean and standard error of mean (SEM). Outliers were tested using ROUT method with the maximum False Discovery Rate (FDR) set to 1%. For the ABR dataset, 4 WT mice (1 male/3 females) and 1 KO (female) mouse were excluded from the analyses because of noisy waveforms, likely caused by physiological artifacts or variability in electrode placements. In a small subset of animals (n = 2 WT and 4 KO) it was not possible to measure the peak ABR wave II amplitude, which is consistent with previous reports of the lower reliability of this wave relative to the others^59,60^. To account for this inconsistency, a linear mixed-effects model by restricted maximum likelihood (REML) was employed to accommodate incomplete repeated-measures for each wave. For the acoustic startle dataset, 1 adult KO male was removed as an outlier, and 1 WT female was removed due to the non-sigmoidal fit. Univariant analysis of variance (x-way ANOVA) was followed by post-hoc multiple comparison tests with two-stage step up method of Benjamini, Krieger and Yekutieli and a discovery threshold of q < 0.05.

## Results

### Altered neuronal responsivity and synchrony in juvenile but not adult Cntnap2 KO mice

We compared ABR thresholds and waveform features in response to broadband clicks between juvenile and adult WT and KO mice across adults and juveniles (**Fig. 1A-B**). ABR measurements from males and females were combined for each genotype, based our previous work in rats that showed an absence of sex differences^34,37^.

First, we compared auditory threshold, which provides an estimate of the auditory sensitivity by identifying the lowest intensity that elicits a reproducible ABR response^54^ (**Fig. 1C**). In *Cntnap2* KO mice, we found a significant effect of age (Two-way ANOVA, F (1, 41) = 4.176, *p* = 0.0475) that indicated greater thresholds in juveniles compared to adult mice, but no effect of genotype, on auditory threshold values (see Table 1; **Fig. 1D**). *Post hoc* analyses revealed no significant differences when comparing between groups. Next, we compared the P2P amplitudes for waveform peaks I-IV (**Fig. 2A**). We were able to reliably detect peaks in all recorded ABR waveforms with the exception of wave II in juveniles: in 2 WT (15.38%) and 4 KO (28.57%), we failed to detect peaks due to their low peak II amplitudes^60^. We found a significant three-way interaction between wave x genotype x age (Three-way ANOVA; F (1.6, 64) = 3.9, *p* = 0.03; **Table 1**) suggesting that the effect of genotype on ABR amplitudes varied across individual waves and differed between juvenile and adult mice. *Post hoc* multiple comparisons tests revealed that juvenile KO mice exhibited significantly larger wave III amplitudes than WT littermates (q = 0.028; **Fig. 2B**). Additionally, we found no significant differences between genotypes in adult mice (q = 0.72), indicating that the difference in wave III amplitude exists only in early development and is normalized in adulthood. Next, we examined full peak width at half-height (FWHH), which represents the degree of neuronal synchronization of firing within each region^61^. We found a significant effect of genotype on FWHH measurements (Three-way ANOVA; F (1, 151) = 4.5, p = 0.04). Interestingly, *post hoc* analyses revealed that juvenile KO mice have narrower peak IV widths than WT littermates (q = 0.002; **Fig. 2C**). Similar to P2P amplitudes, we did not detect any differences between adult WT and KO mice, suggesting that the phenotype normalizes through development.

**Table 1.**
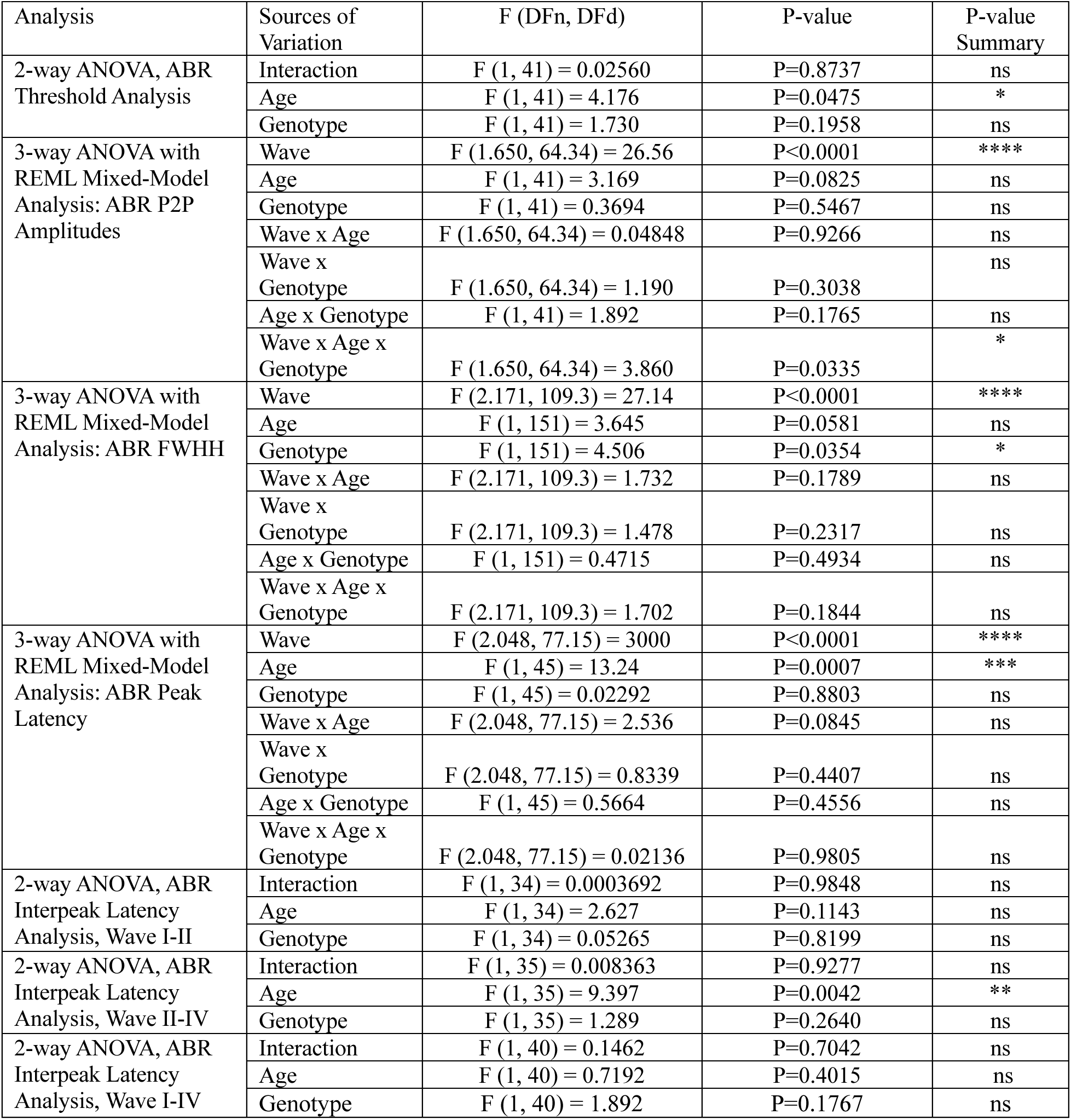
ABR statistics.

| Analysis | Sources of Variation | F (DFn, DFd) | P-value | P-value Summary |
| --- | --- | --- | --- | --- |
| 2-way ANOVA, ABR Threshold Analysis | Interaction | F (1, 41) = 0.02560 | P=0.8737 | ns |
|  | Age | F (1, 41) = 4.176 | P=0.0475 | * |
|  | Genotype | F (1, 41) = 1.730 | P=0.1958 | ns |
| 3-way ANOVA with REML Mixed-Model Analysis: ABR P2P Amplitudes | Wave | F (1.650, 64.34) = 26.56 | P<0.0001 | **** |
|  | Age | F (1, 41) = 3.169 | P=0.0825 | ns |
|  | Genotype | F (1, 41) = 0.3694 | P=0.5467 | ns |
|  | Wave x Age | F (1.650, 64.34) = 0.04848 | P=0.9266 | ns |
|  | Wave x Genotype | F (1.650, 64.34) = 1.190 | P=0.3038 | ns |
|  | Age x Genotype | F (1, 41) = 1.892 | P=0.1765 | ns |
|  | Wave x Age x Genotype | F (1.650, 64.34) = 3.860 | P=0.0335 | * |
| 3-way ANOVA with REML Mixed-Model Analysis: ABR FWHH | Wave | F (2.171, 109.3) = 27.14 | P<0.0001 | **** |
|  | Age | F (1, 151) = 3.645 | P=0.0581 | ns |
|  | Genotype | F (1, 151) = 4.506 | P=0.0354 | * |
|  | Wave x Age | F (2.171, 109.3) = 1.732 | P=0.1789 | ns |
|  | Wave x Genotype | F (2.171, 109.3) = 1.478 | P=0.2317 | ns |
|  | Age x Genotype | F (1, 151) = 0.4715 | P=0.4934 | ns |
|  | Wave x Age x Genotype | F (2.171, 109.3) = 1.702 | P=0.1844 | ns |
| 3-way ANOVA with REML Mixed-Model Analysis: ABR Peak Latency | Wave | F (2.048, 77.15) = 3000 | P<0.0001 | **** |
|  | Age | F (1, 45) = 13.24 | P=0.0007 | *** |
|  | Genotype | F (1, 45) = 0.02292 | P=0.8803 | ns |
|  | Wave x Age | F (2.048, 77.15) = 2.536 | P=0.0845 | ns |
|  | Wave x Genotype | F (2.048, 77.15) = 0.8339 | P=0.4407 | ns |
|  | Age x Genotype | F (1, 45) = 0.5664 | P=0.4556 | ns |
|  | Wave x Age x Genotype | F (2.048, 77.15) = 0.02136 | P=0.9805 | ns |
| 2-way ANOVA, ABR Interpeak Latency Analysis, Wave I-II | Interaction | F (1, 34) = 0.0003692 | P=0.9848 | ns |
|  | Age | F (1, 34) = 2.627 | P=0.1143 | ns |
|  | Genotype | F (1, 34) = 0.05265 | P=0.8199 | ns |
| 2-way ANOVA, ABR Interpeak Latency Analysis, Wave II-IV | Interaction | F (1, 35) = 0.008363 | P=0.9277 | ns |
|  | Age | F (1, 35) = 9.397 | P=0.0042 | ** |
|  | Genotype | F (1, 35) = 1.289 | P=0.2640 | ns |
| 2-way ANOVA, ABR Interpeak Latency Analysis, Wave I-IV | Interaction | F (1, 40) = 0.1462 | P=0.7042 | ns |
|  | Age | F (1, 40) = 0.7192 | P=0.4015 | ns |
|  | Genotype | F (1, 40) = 1.892 | P=0.1767 | ns |

To assess whether the speed of auditory processing in the early ascending auditory brainstem pathway was altered due to the lack of the *Cntnap2* gene, we compared peak latencies between WT and KO mice across the two age groups. As expected from previous reports^34,62^, there was a significant effect of age on wave latencies (Three-way ANOVA; F (1, 45) = 13.24, *p* < 0.001, **Table 1**), where adult WT mice displayed significantly lower latencies than juvenile WT mice (*post hoc* comparisons: q = 0.046) for wave I. Adult KO mice also exhibited shorter average latency value than juvenile KO mice, although this difference did not reach statistical significance (q = 0.069). Additionally, we calculated interpeak latencies by subtracting the peak latencies of the respective peaks (i.e. WII-IV = wave IV – wave II) to further dissect whether conduction time is altered in specific regions of the auditory pathway. We found an age effect for WII-IV interpeak latency (Two-way ANOVA; F (1, 35) = 9.397, *p* < 0.01, **Table 1**) but not for WI-II, WII-IV, or WI-IV interpeak latencies (**Fig. 2D**). *Post hoc* testing revealed no significant pairwise comparisons for each of the interpeak latencies (**Table 1**).

In summary, juvenile *Cntnap2* KO mice exhibit higher wave III peak amplitudes and narrower wave IV peak widths, waveform characteristics that respectively represent the summated neural activity in the superior olivary complex (SOC) and the inferior colliculus (IC), two brainstem nuclei involved in early auditory processing^63^. Additionally, both phenotypes are normalized in adult mice, showing that these ABR phenotypes are developmentally transient. Surprisingly, we did not detect any genotype differences in latency across all waves in both juvenile and adult mice, highlighting that the speed of auditory processing within the *Cntnap2* KO mouse model may be conserved.

### Juvenile Cntnap2 KO mice exhibit exaggerated startle that persists into adulthood in a sex-specific manner

Since the above ABR results revealed altered brainstem processing of auditory information within juvenile KO mice, we next asked whether these changes were accompanied by alterations in the acoustic startle behavioural responses. Although ABRs and the acoustic startle response assess difference aspects of auditory function, they are functionally related through their shared dependence on auditory neural encoding within the auditory brainstem^64^. Thus, alterations in auditory processing can modify the strength of acoustic input reaching the startle circuit, contributing to potential differences in the magnitude of the behavioural response^65,66^. To test this, we assessed the acoustic startle response in juvenile and adult WT and KO mice across a range of stimulus intensities (**Fig. 3A**). First, to rule out the contributions of the animals’ body mass on startle magnitude, we examined whether the maximum startle response correlated with the weight of each animal. We did not find any significant correlations in both male and female juvenile mice regardless of genotype (**Fig. 3B**; WT Male, R^2^ = 0.16, F (1,9) = 1.7, *p* = 0.23; WT Female, R^2^ = 0.04, F (1,7) = 0.30, *p* = 0.60; KO Male, R^2^ = 0.1, F (1,8) = 1.0, *p* = 0.34; KO Female, R^2^ = 0.01, F (1,9) = 0.08, *p* = 0.78). Next, we compared the maximum startle magnitude of juvenile and adult *Cntnap2* KO mice to their age-matched WT littermates. In juveniles, we found a significant effect of genotype (Two-way ANOVA, F (1,35) = 18.1, *p* < 0.0001) on maximum startle response (**Fig. 3C,E**). *Post hoc* comparisons found that both juvenile male and female KO mice had larger maximum startle responses than their age-matched WT littermates (Male q = 0.04, Female q = 0.01; **Fig. 3E**), indicating an exaggerated startle response in this age group. In contrast, we did not detect any differences in threshold, ES50, and saturation between juvenile WT and KO mice, in both sexes (**Fig. 3D, F-H**; **Table 2**)

**Table 2.** Startle and PPI statistics.

| Analysis | Sources of Variation | F (DFn, DFd) | P-value | P-value Summary |
| --- | --- | --- | --- | --- |
| 2-way ANOVA, Juvenile Maximum Startle Response (*Top*) | Interaction | F (1, 41) = 0.03399 | P=0.8546 | ns |
|  | Sex | F (1, 41) = 0.5272 | P=0.4719 | ns |
|  | Genotype | F (1, 41) = 18.08 | P=0.0001 | *** |
| 2-way ANOVA, Juvenile Startle Threshold | Interaction | F (1, 42) = 0.9207 | P=0.3428 | ns |
|  | Sex | F (1, 42) = 0.2800 | P=0.5995 | ns |
|  | Genotype | F (1, 42) = 1.705 | P=0.1987 | ns |
| 2-way ANOVA, Juvenile Startle ES50 | Interaction | F (1, 42) = 0.01569 | P=0.9009 | ns |
|  | Sex | F (1, 42) = 1.848 | P=0.1812 | ns |
|  | Genotype | F (1, 42) = 0.2557 | P=0.6157 | ns |
| 2-way ANOVA, Juvenile Startle Saturation Point | Interaction | F (1, 42) = 0.9619 | P=0.3323 | ns |
|  | Sex | F (1, 42) = 1.973 | P=0.1675 | ns |
|  | Genotype | F (1, 42) = 0.1904 | P=0.6648 | ns |
| 2-way ANOVA, Adult Maximum Startle Response (*Top*) | Interaction | F (1, 39) = 1.416 | P=0.2413 | ns |
|  | Sex | F (1, 39) = 8.549 | P=0.0057 | ** |
|  | Genotype | F (1, 39) = 4.502 | P=0.0403 | * |
| 2-way ANOVA, Adult Startle Threshold | Interaction | F (1, 39) = 0.3691 | P=0.5470 | ns |
|  | Sex | F (1, 39) = 2.970 | P=0.0927 | ns |
|  | Genotype | F (1, 39) = 1.948 | P=0.1707 | ns |
| 2-way ANOVA, Adult Startle ES50 | Interaction | F (1, 39) = 2.953 | P=0.0936 | ns |
|  | Sex | F (1, 39) = 4.324 | P=0.0442 | ns |
|  | Genotype | F (1, 39) = 0.4414 | P=0.5104 | ns |
| 2-way ANOVA, Adult Startle Saturation Point | Interaction | F (1, 39) = 3.118 | P=0.0853 | ns |
|  | Sex | F (1, 39) = 1.913 | P=0.1745 | ns |
|  | Genotype | F (1, 39) = 0.05485 | P=0.8161 | ns |
| 2-way ANOVA, Juvenile PPI (75 dB prepulse, 30 ms ISI) | Interaction | F (1, 42) = 0.5989 | P=0.4433 | ns |
|  | Sex | F (1, 42) = 0.2644 | P=0.6098 | ns |
|  | Genotype | F (1, 42) = 1.863 | P=0.1795 | ns |
| 2-way ANOVA, Juvenile PPI (75 dB prepulse, 100 ms ISI) | Interaction | F (1, 42) = 0.6811 | P=0.4139 | ns |
|  | Sex | F (1, 42) = 0.0006977 | P=0.9791 | ns |
|  | Genotype | F (1, 42) = 1.837 | P=0.1826 | ns |
| 2-way ANOVA, Juvenile PPI (85 dB prepulse, 30 ms ISI) | Interaction | F (1, 42) = 1.954 | P=0.1695 | ns |
|  | Sex | F (1, 42) = 0.06245 | P=0.8039 | ns |
|  | Genotype | F (1, 42) = 0.5579 | P=0.4593 | ns |
| 2-way ANOVA, Juvenile PPI (85 dB prepulse, 100 ms ISI) | Interaction | F (1, 42) = 0.2056 | P=0.6526 | ns |
|  | Sex | F (1, 42) = 1.096 | P=0.3011 | ns |
|  | Genotype | F (1, 42) = 0.04927 | P=0.8254 | ns |
| 2-way ANOVA, Adult PPI (75 dB prepulse, 30 ms ISI) | Interaction | F (1, 43) = 6.171 | P=0.0170 | * |
|  | Sex | F (1, 43) = 4.550 | P=0.0387 | * |
|  | Genotype | F (1, 43) = 2.814 | P=0.1007 | ns |
| 2-way ANOVA, Juvenile PPI (75 dB prepulse, 100 ms ISI) | Interaction | F (1, 43) = 4.893 | P=0.0323 | * |
|  | Sex | F (1, 43) = 1.857 | P=0.1800 | ns |
|  | Genotype | F (1, 43) = 0.1813 | P=0.6724 | ns |
| 2-way ANOVA, Juvenile PPI (85 dB prepulse, 30 ms ISI) | Interaction | F (1, 43) = 4.666 | P=0.0364 | * |
|  | Sex | F (1, 43) = 0.2245 | P=0.6380 | ns |
|  | Genotype | F (1, 43) = 5.964e-005 | P=0.9939 | ns |
| 2-way ANOVA, Juvenile PPI (85 dB prepulse, 100 ms ISI) | Interaction | F (1, 43) = 3.867 | P=0.0557 | ns |
|  | Sex | F (1, 43) = 1.832 | P=0.1829 | ns |
|  | Genotype | F (1, 43) = 0.3570 | P=0.5533 | ns |

Next, we compared the startle response in adult WT and KO mice. Again, we examined the relationship between the weight of mice and their maximum startle magnitude in the adult group and did not find any significant correlations in both male and female adult mice regardless of genotype (**Fig. 4A**, WT Male, R^2^ = 0.01, F(1,10) = 0.05, *p* = 0.83; WT Female, R^2^ = 0.06, F(1,8) = 0.53, *p* = 0.48; KO Male, R^2^ = 0.01, F(1,8) = 0.05, *p* = 0.82; KO Female, R^2^ = 0.01, F(1,10) = 0.12, *p* = 0.73). Similarly to juveniles, we observed both a significant genotype difference in the maximum startle response of adult mice (F (1, 39) = 4.5, *p* = 0.04), as well as a sex difference (Two-way ANOVA, F (1, 39) = 8.5, *p* = 0.01; **Fig. 4B,D**). Interestingly, *post hoc* analyses revealed that adult male KO mice startled at greater magnitudes compared to male WT mice (q = 0.04), and female KO mice (q = 0.01), while female WT and KO mice exhibited similar startle magnitudes (q = 0.39; **Fig. 4D**). Given our above data showing higher startle magnitudes in juvenile KO mice of both sexes (**Fig. 3**), these results suggest that the exaggerated startle phenotype persists into adulthood in male KO mice, whereas the phenotype becomes normalized in adult female KO mice. We also observed a significant effect of sex on ES50 (Two-way ANOVA, F (1, 39) = 4.324, *p* = 0.04; **Fig. 4C,F**) in which males appeared to have higher values, but no significant *post hoc* comparisons were found (WT q = 0.07; KO q = 0.83). We did not detect any differences in threshold and saturation between juvenile WT and KO mice, in both sexes (**Fig. 4C,E,H**; **Table 2**)

**Figure 4.**
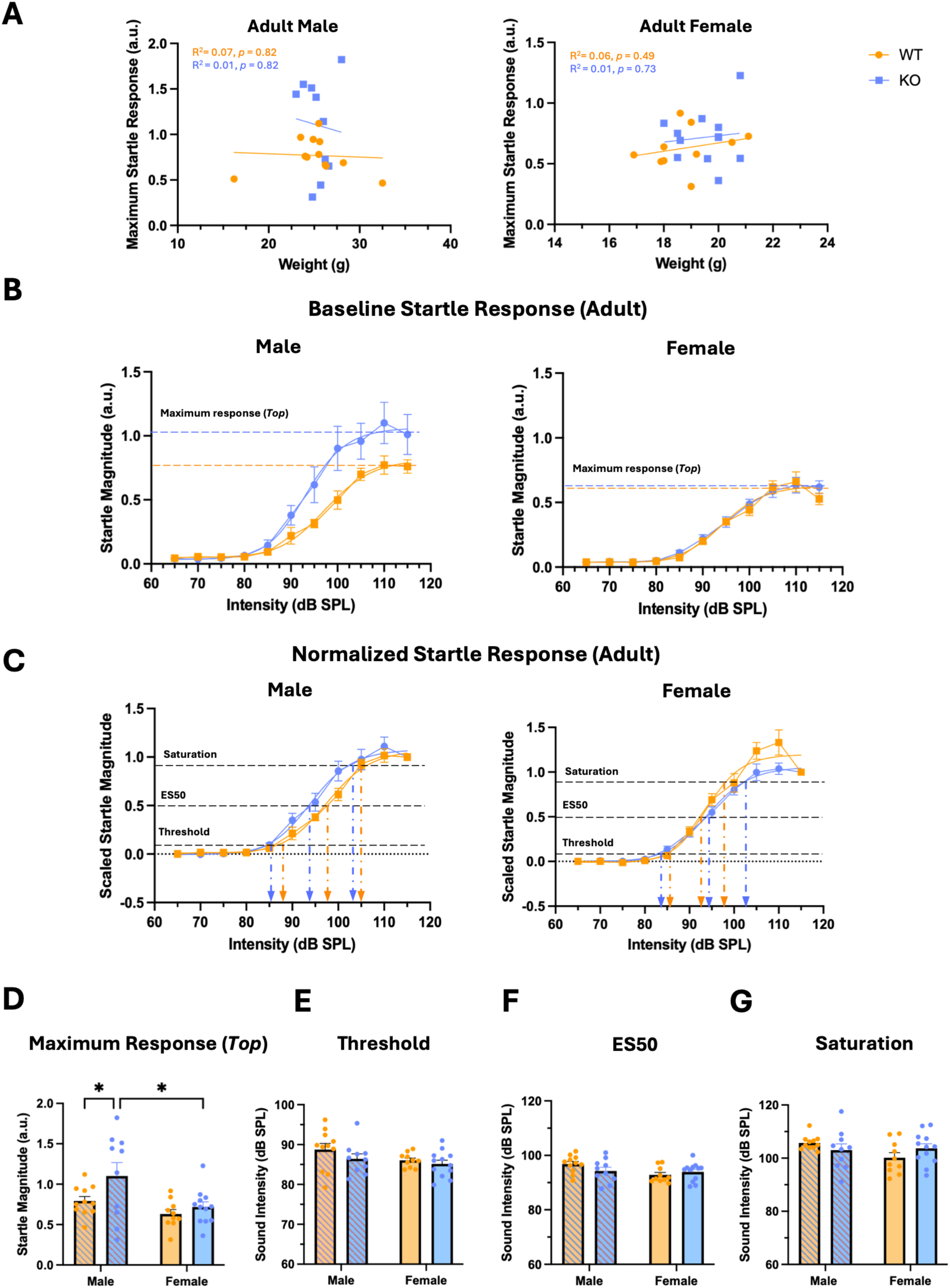
Adult male *Cntnap2* KO mice display a persistent, exaggerated startle response. Bar graphs show individual values with error bars representing SEM. **(A)** Correlation between weight and maximum startle response for each sex. Shown are I/O functions that represent the mean and standard error of mean (SEM) for each intensity with fitted sigmoidal curve for **(B)** Adult baseline startle response curves separated by sex**. (C)** Scaled startle response curve for each sex. Bar graphs show the mean, individual values, and error bars representing SEM for **(D)** Maximum startle response (*Top*), **(E)** Startle thresholds, **(F)** ES50, and **(G)** saturation points separated by sex. \**p* < 0.05.

Based on our previous findings in rats^34,37,39,67^, we predicted that, in addition to startle reactivity, sensorimotor gating would also be affected in *Cntnap2* KO mice during development and in adulthood. Therefore, we compared PPI between WT and KO mice, at juvenile and adult stages. To test for PPI, we presented different conditions of prepulses that varied in intensity (75- or 85-dB SPL) and ISI (30 or 100 ms) to test the sensitivity of the sensorimotor gating system under these contexts^34^ (**Fig. 5A**). In juvenile mice, we did not detect any significant differences between genotypes or sex, nor interactions between the two, in any of the PPI conditions for either age groups (see Table 2, **Fig. 5B**). However, in adults, there was a significant interaction between genotype and sex (Two-wav ANOVA, F (1, 41) = 5.833, *p* = 0.02, **Fig. 5C**) when a 75 dB prepulse was presented 30 ms before the startle stimulus. *Post hoc* analyses revealed that female WT mice have a lower percentage of PPI compared to both WT male (q = 0.01) and KO females (q = 0.01). Additionally, there was also an interaction between genotype and sex (Two-way ANOVA, F (1, 41) = 5.399, *p* = 0.03) when a 75 dB prepulse was presented 100 ms before the startle stimulus, but no differences were detected between specific groups following *post hoc* corrections (**Table 2**).

**Figure 5.**
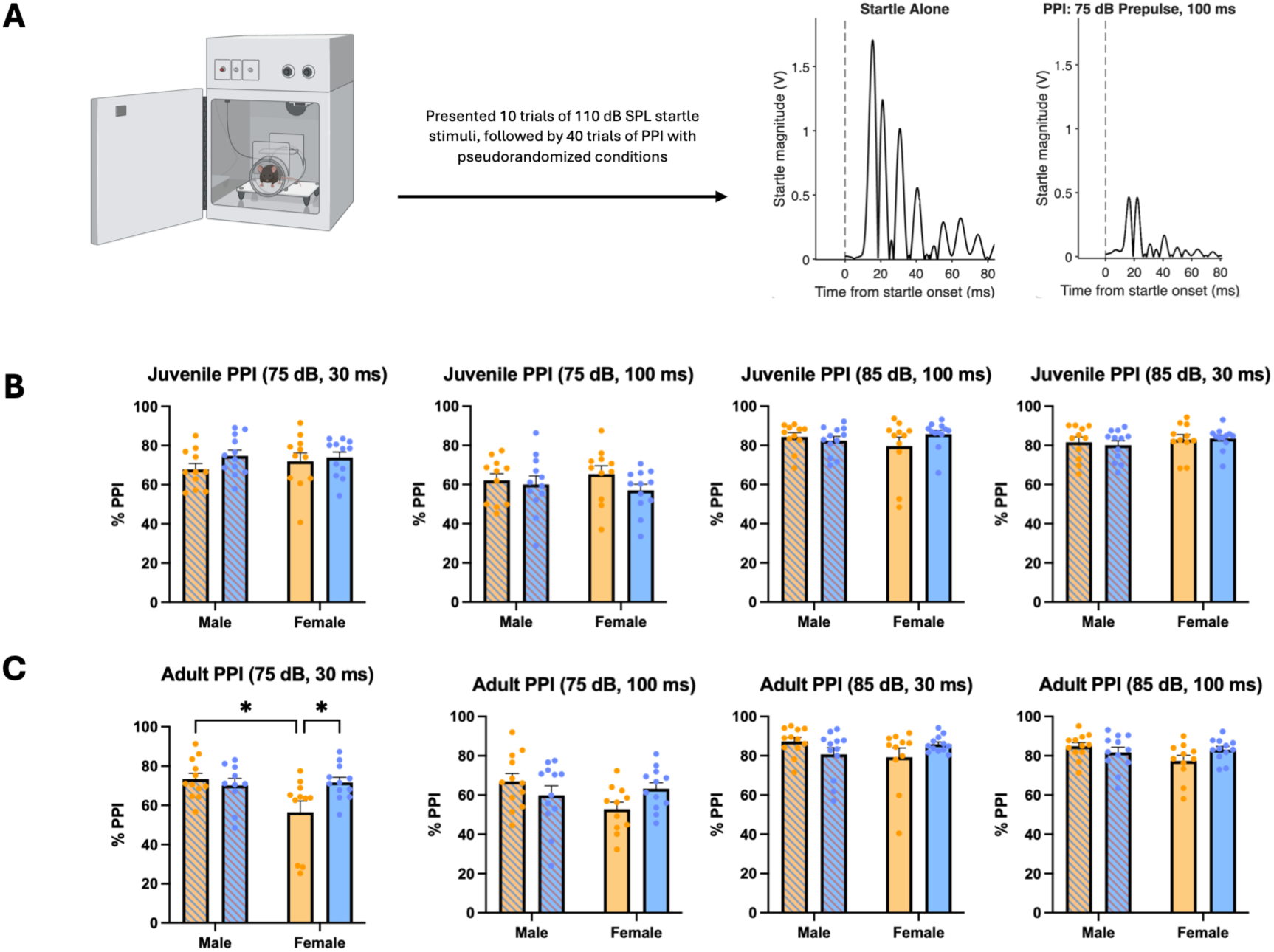
Sensorimotor gating is generally preserved in both juvenile and adult *Cntnap2* KO mice. **(A)** PPI Experimental timeline with example traces of startle alone and the 75 dB prepulse, 100ms PPI condition. Bar graphs compare WT (yellow) and KO (blue) % PPI responses. Individual values are represented as dots, and error bars represent SEM for **(B)** juvenile mice **(C)** adult mice across the four different conditions – 75 dB or 85 dB prepulse at an interstimulus intervals of 30 or 100 ms. \**p* < 0.05.

Taken together, these results show that although juvenile KO mice of both sexes and adult male KO mice display exaggerated startle responses, their PPI responses remain largely intact. Interestingly, adult KO females exhibit greater PPI compared to age-matched WT littermates at lower-intensity prepulses, potentially revealing a sex-specific alteration in sensorimotor gating that emerges in adulthood.

## Discussion

In this study, we examined how disruptions in the *CNTNAP2* gene affect auditory brainstem processing by characterizing ABR morphology and startle reactivity using the *Cntnap2* KO mouse model. We found that juvenile KO mice exhibit higher wave III peak amplitude and narrower wave IV peak width in response to auditory stimuli. Both phenotypes were normalized in adulthood, suggesting developmentally transient alterations in neural responsivity and synchrony within discrete auditory brainstem circuits. Behaviourally, juvenile KO mice exhibited an exaggerated startle response that normalized in adult females but persisted in adult males. We also identified a sex-specific alteration in PPI, with adult KO females exhibited higher PPI compared to adult WT females. These results reveal that, despite the developmentally transient alteration of auditory responsivity in the early auditory pathway, auditory startle phenotypes manifest into adulthood in a sex-specific manner, suggesting persistent changes in the behavioural circuitry that are influenced by sex.

Our results further our understanding of the role of *CNTNAP2* in auditory processing, extending auditory phenotypes previously reported in humans^68^ and rats^34,35,37,39,69^. Our finding that wave III peak amplitudes are higher in juvenile KO mice is consistent with previous reports in *Cntnap2* KO rats^34^. The waveform characteristics of wave III reflect the neural activity responses that originate from the SOC which is the first major brainstem region that receives binaural input^70^. Our observation of increased wave III amplitude phenotype suggests that the circuitry within the SOC may be hyper-responsive to incoming auditory inputs. This is in line with a previous report from *Fmr1* KO mice, another loss-of-function genetic model for ASD^71^, which also display increased ABR wave III amplitudes that correspond with hypersensitivity to sounds, as well as higher firing rates in the lateral superior olive (LSO) of the SOC driven by excessive excitatory inputs^72^. Given the known role of the LSO in integrating excitatory and inhibitory signals for binaural auditory processing^72,73^, the enhanced wave III amplitudes in juvenile *Cntnap2* KO mice may reflect disruption in the balance between excitatory and inhibitory synaptic inputs, which may alter the developmental refinement of the SOC circuitry to cause increased neural responsivity to sound.

Another ABR phenotype we found in juvenile KO mice is a narrower wave IV peak. Given that wave IV reflects neural responses in the IC^63,74^, these results may indicate a more synchronous neuronal firing in the IC of KO mice in response to auditory stimuli, consistent with IC *in vivo* recording results from juvenile *Fmr1* KO mice^72^. Although *Cntnap2* and *Fmr1* are two genes that encode for protein products with distinct molecular functions^25,75^, loss-of-function mutation of these genes both cause ASD-associated phenotypes in rodent model including atypical auditory processing^34,72^ as well as many other neurobiological phenotypes^51,76–79^. Thus, these shared electrophysiological phenotypes within the IC may be a convergent contributing mechanism underlying atypical auditory processing in these two well-established genetic mouse models of ASD. Future studies should examine the firing properties and morphology of the SOC and IC to further explore how disruptions in the refinement of brainstem circuitry can underlie auditory processing disruptions in the *Cntnap2* KO mice.

Contrary to the findings from *Cntnap2* KO rats and humans, we did not observe prolonged latencies across any of the ABR waveforms^8,34,62^. These differences may reflect species-specific organization of the auditory brainstem that result in distinct compensatory mechanisms^80^. Mice also have shorter and more compact auditory brainstems, which can reduce the sensitivity of ABR latency measurements^81^. Additionally, following auditory perturbation, mice are more resilient to peripheral auditory damage and exhibit increased brainstem gain to preserve central function compared to rats^80^.

Thus, these opposing central gain regulations may shift where *Cntnap2*-related phenotypes manifest along the auditory pathway, potentially preserving auditory information transmission within the brainstem intact in mice.

Despite the normalization of ABR phenotypes in adulthood, we found that adult KO males show exaggerated startle responses that persisted into adulthood while adult KO females exhibit normalized startle responses, findings which are mainly in accordance with the findings in rats. Auditory startle is regulated by a brainstem pathway involving spiral ganglion cells in the auditory nerve synapsing onto cochlear root neurons that innervate the pontine reticular nucleus (PnC) which send ipsilateral projections to motor neurons within the spinal cord^69^. The PnC integrates convergent excitatory and inhibitory inputs^82^, many of which originate from brain regions known to exhibit sex-dependent differences in connectivity and hormonal regulation^83,84^. Consistent with this, male and female *Cntnap2* KO rats exhibit distinct PnC neural properties that are linked to an exaggerated startle response^40^. In the present study, the exaggerated startle response, observed predominantly in male *Cntnap2* KO mice, may reflect sex differences in the regulation or developmental compensation of atypical inhibitory signaling within the startle pathway. This may result in greater PnC excitability and enhanced auditory sensitivity, as evidenced by the leftward shift in the startle I/O function in male *Cntnap2* KO rats^37,67^. In contrast, females may recruit compensatory mechanisms that preserve the excitation-inhibition balance to limit the persistence of startle abnormalities^85,86^.

Interestingly, the PPI responses were generally comparable between WT and KO mice, at both juvenile and adult stages. No sex differences were detected, except for adult female WT mice at the 75 dB prepulse, 30 ms interstimulus interval condition, which exhibited lower % PPI compared to WT males as well as KO males and females. In both human and rodent studies, females have been reported to have lower PPI compared to males^87^. Previous reports have found that *Cntnap2* KO mice have intact PPI^53,88^, while data from *Cntnap2* KO rats suggest lower PPI when prepulse was at a higher intensity^34^. One possible explanation for this difference is that, in our study, PPI was assessed at only one startle intensity (110 dB) which may not have been sufficiently sensitive to detect subtle group differences^37^. Previous studies have shown that PPI magnitude is dependent on both prepulse and startle pulse conditions, especially if baseline startle response magnitudes are different between experimental groups as in our case^37,39^. Future studies assessing PPI across multiple startle pulse intensities and accounting for differences in baseline startle would help provide a more comprehensive assessment of sensorimotor gating mechanisms.

By utilizing both physiological and behavioural assessments of auditory brainstem function, we show that *Cntnap2* KO mice display abnormalities early in development that may lead to persistent changes in downstream behavioural circuits that rely on auditory inputs. Overall, our findings reveal that the loss of *CNTNAP2* disrupts early auditory processing during development in mice similarly to what has been reported in rats, with some distinct elements only observed in mice. Consistent with findings from both rat and human studies, our results suggest that alterations in auditory brainstem processing are associated with persistent changes in auditory-evoked behavioural responses, even as ABR abnormalities normalize with maturation. Notably, our findings further reveal a sex-dependent persistence of this behavioural phenotype, as the exaggerated startle was observed selectively in adult male KO mice.

Together, these findings highlight the important role of *Cntnap2* in the developing auditory brainstem, and that early sensory processing abnormalities caused by a loss of *Cntnap2,* although transient, may contribute to the development of more complex behavioural phenotypes that persist through adulthood. Finally, our finding of sex differences in the developmental correction of early sensory processing phenotypes emphasizes the consideration of sex as a critical biological variable when identifying mechanisms and potential targets for early interventions in ASD.

## Acknowledgments

We acknowledge members of the Choe Lab help with discussion and feedback. We also acknowledge K. Hossain, Y. K. Lee, and undergraduate research assistants in the Choe lab for mouse colony management support and genotyping. Finally, we thank the Central Animal Facility (CAF) staff, Dr. Tim Ryan, and Dr. Michelle Reichart for their continued animal care support.

